# Sarcomere length, fascicle length, and serial sarcomere number are preserved in paretic hindlimb muscles following chronic stroke in rats despite persistent motor impairment

**DOI:** 10.64898/2026.09.09.750457

**Authors:** Stephanie A. Ross, Sydney N. Dorscher, Timothy R. Leonard, Ruth A. Seerattan, Dale Corbett, Walter Herzog

## Abstract

Stroke causes motor impairments that are commonly attributed to altered neural control, but chronic changes in neural activation and muscle use may also influence skeletal muscle structure. This study examined whether chronic stroke alters sarcomere length and dispersion, fascicle length, or serial sarcomere number in adult skeletal muscle. Twenty-four female Sprague-Dawley rats underwent photothrombotic stroke or sham surgery. Limb-specific motor impairment was assessed longitudinally using a beam traversal task. At 4.5 months post-surgery, the lateral gastrocnemius and extensor digitorum longus muscles were harvested from paretic and non-paretic limbs. We measured fascicle length directly from isolated fascicle bundles and quantified sarcomere length and dispersion using laser diffraction. Stroke animals exhibited persistent impairment of the paretic limb during beam traversal with elevated misstep rates. Despite this persistent motor impairment, sarcomere length and dispersion did not differ between stroke and sham animals or between limbs. Fascicle length and serial sarcomere number were greater in stroke than sham animals, but these differences were not limb-specific and were therefore unlikely to reflect stroke-related changes. Fascicle length, sarcomere length and dispersion, and serial sarcomere number differed between the lateral gastrocnemius and extensor digitorum longus, consistent with differences in architecture or relative length at the selected joint angles. These findings indicate that persistent neural impairment following adult-onset stroke does not necessarily result in substantial changes in sarcomere length or sarcomeres in series. Instead, sarcomere changes may depend on additional factors, including the timing of neural injury relative to growth and the mechanical environment experienced by the muscle.

## Introduction

Stroke is a leading cause of long-term motor disability (Mendis, 2013), with persistent impairments in strength, coordination, and locomotor function that can limit independence and quality of life (Mayo et al., 1999; Patel et al., 2006). These impairments are commonly attributed to changes in the central nervous system that alter motor control, including inappropriate activation, abnormal muscle synergies, and impaired coordination of movement (Arene and Hidler, 2009; Li et al., 2021). However, skeletal muscle is not simply a passive effector of neural commands. Muscle is a highly adaptable tissue that remodels in response to changes in neural activation (Wilson and Deschenes, 2005), mechanical loading (Wisdom et al., 2015), and muscle length (Blazevich et al., 2025). Because stroke produces persistent changes in neural input and muscle use, it may also induce structural and functional adaptations within skeletal muscle that could contribute to or exacerbate motor deficits.

Evidence from other neurological conditions supports the possibility that altered neural and mechanical environments can influence the longitudinal structure of skeletal muscle. Cerebral palsy (CP), a neurological condition associated with early-life brain injury and persistent motor impairment, is characterized by alterations in skeletal muscle structure, including increased sarcomere length and fewer sarcomeres in series (Leonard et al., 2019; Lieber and Fridén, 2002; Mathewson et al., 2015; Smith et al., 2011). These changes have been proposed to result, at least in part, from impaired sarcomerogenesis during muscle growth, such that muscles fail to add sufficient sarcomeres in series as the skeleton increases in length. Because this process occurs during development, it is unclear whether the longitudinal muscle adaptations observed in CP would also occur following neurological injury after skeletal muscle maturation. Nevertheless, the CP literature demonstrates that chronic alterations in neural activation and the mechanical environment can be associated with remodelling of sarcomere organization. This raises the question of whether persistent neural impairment following adult-onset stroke is sufficient to produce similar adaptations in mature skeletal muscle.

Evidence regarding longitudinal muscle structure following stroke is limited. In individuals with chronic stroke, Adkins et al. (2021) reported shorter fascicles and fewer sarcomeres in series in the paretic compared with the non-paretic biceps brachii, but no difference in mean sarcomere length. These findings suggest that fascicle length and the number of sarcomeres arranged in series may be altered following stroke without a corresponding change in mean sarcomere length. However, sarcomere length is heterogeneous within muscle, and localized measurements may not necessarily represent the sarcomere length throughout an entire fascicle. In addition, fascicle length measurements obtained using ultrasound require visualization or reconstruction of the fascicle trajectory and may be affected by muscle composition, imaging quality, and the passive mechanical conditions under which the muscle is assessed. These limitations make it difficult to determine whether the reported differences reflect true remodelling of longitudinal muscle structure. Furthermore, whether chronic stroke alters sarcomere length dispersion, which may provide information about changes in sarcomere organization that are not captured by mean sarcomere length, remains unclear.

Direct measurements of sarcomere and fascicle length in an animal model provide an opportunity to determine whether chronic neural impairment is accompanied by remodelling of longitudinal muscle structure under controlled experimental conditions. Therefore, the purpose of this study was therefore to determine whether chronic stroke alters sarcomere length, sarcomere length dispersion, fascicle length, or serial sarcomere number in the lateral gastrocnemius and extensor digitorum longus muscles of rats. Dissected fascicle bundles were isolated from each muscle, fascicle length was measured directly, and laser diffraction was used to determine sarcomere length and sarcomere length dispersion. Serial sarcomere number was calculated from measured fascicle length and sarcomere length. These properties were compared between paretic and non-paretic limbs of stroke animals and between corresponding limbs of stroke and sham animals. Motor function was assessed longitudinally using a beam traversal task to establish the persistence of motor impairment throughout the post-stroke period.

## Materials and Methods

### Animals

Twenty-four 14-week-old female Sprague-Dawley rats from the University of Calgary on-campus colony were included in the study (body mass at time of surgery (mean ± s.d.): 278.0 ± 17.5 g). Animals were randomly allocated to either a stroke group (n = 17) or sham stroke group (n = 7). Rats were housed two per cage under a conventional 12-hour light/12-hour dark cycle, with food and water provided ad libitum. Cages did not contain running wheels, and animals were housed in a room with sentinel animals for health monitoring. Body mass was measured immediately prior to stroke or sham stroke surgery and again immediately prior to euthanasia. Husbandry and veterinary care were provided and overseen by the University of Calgary Life and Environmental Sciences Animal Research Centre. All procedures were approved by the University of Calgary Animal Care Committee (protocol AC22-0198).

### Stroke and sham stroke induction

Stroke was induced using photothrombosis. We administered analgesics (subcutaneous buprenorphine, 0.1 mg/kg, and meloxicam, 2 mg/kg) and sterile saline (10 mg/kg) prior to surgery, then anaesthetized the animals with inhaled isoflurane (5% induction, 2% maintenance in 1 L/min O_2_). Once the rats were deeply anaesthetized, we shaved the scalp and positioned the animal in a stereotaxic frame, securing the skull position with ear bars. We then made a midline skin incision over the skull, followed by blunt dissection and removal of the periosteum to expose the skull surface. We positioned an aluminum foil aperture (5 mm mediolateral x 7 mm anteroposterior; Jeffers et al. (2020)) over the hindlimb sensorimotor cortex (−1.5 mm anterior- and ±3 mm medial-lateral relative to Bregma; McDonald et al. (2021)), with the hemisphere selected for stroke induction randomized across animals to control for potential effects of limb dominance. We then positioned a light source (Fiber-Lite MI-LED B1 high-intensity LED illuminator; Dolan-Jenner, Boxborough, MA, USA) over the aperture and injected a photoactive dye (Rose Bengal; MilliporeSigma Canada Ltd., Oakville, ON, Canada; St. 20 mg/kg) in sterile saline via the lateral tail vein and allowed 2 minutes for the dye to circulate systemically. We then activated the light source for 20 minutes, a duration determined necessary for successful stroke induction for our light source during pilot trials. After illumination, we turned off the light, closed the skin incision, and removed the animal from the stereotaxic frame before discontinuing the anesthesia. Postoperative analgesia was provided for 24 hours following surgery (buprenorphine, 0.1 mg/kg every 6-12 hours; meloxicam, 2 mg/kg every 24 hours). For the sham group, we followed the same surgical procedures except that the light source was not turned on. Unilateral stroke induction resulted in hemiparesis of the contralateral limb; we designated the affected limb contralateral to the lesion as the paretic limb and the limb on the opposite side as the non-paretic limb.

#### Beam traversal task to quantify limb-specific motor impairments

We used a tapered beam traversal task (Schallert et al., 2002) to detect impairments in limb placement following stroke. The task required animals to traverse a tapered beam from its wider end to a goal box at the opposite end, requiring precise foot placement to maintain balance while walking across the beam. Each animal underwent training twice daily, 5 days per week for 2 weeks, completing six trials during each session to traverse the beam before baseline trials were conducted prior to stroke or sham stroke surgery. Following surgery, we tested animals weekly for six beam trials per animal. Trials were video recorded and manually analyzed by an investigator blinded to group assignment to determine the number of missteps per limb, defined as the number of times a foot fell off the side of the beam divided by the total number of steps taken by that limb. We calculated the proportion of missteps separately for each limb and averaged the values across the six trials to obtain a weekly measure of beam traversal performance per limb. To assess changes in performance over the recovery period, values from weeks 1 and 2 were averaged to represent early post-stroke performance, and values from weeks 16 and 17 were averaged to represent late post-stroke performance.

### Euthanasia, harvesting, and brain histology

At 18 weeks (4.5 months) post-surgery, animals were euthanized via thoracotomy while deeply anesthetized (5% inhaled isoflurane in 1 L/min O_2_). Whole brains were carefully removed from the skull, weighed, and the cerebellum was removed prior to fixation in neutral buffered formalin (NBF) at room temperature overnight. The following day, brains were placed in a brain matrix to standardize subsequent sectioning. Brains were then sectioned into three pieces, with cuts made approximately 3-4 mm from the frontal end and 5-6 mm from the posterior end to obtain a 6-8 mm section encompassing the region of the lesion. The resulting brain sections were placed in fresh NBF overnight at 4°C. Tissue was then cryoprotected by immersion in 10%, 20%, and 30% sucrose phosphate buffer solutions over 4 days at 4°C, with the solution changed daily. Sections were subsequently embedded in molds with optimal cutting temperature (OCT) compound and frozen in liquid nitrogen. Thirty-micrometer sections were collected from the middle brain segment, beginning at the frontal end, and mounted onto Fisherbrand Superfrost Plus slides. Slides were dried on a warm plate overnight before staining with freshly prepared Nissl stain (0.1% cresyl violet in distilled water). Briefly, sections were defatted in a 1:1 solution of alcohol and chloroform overnight and then stained in cresyl violet solution for 10 min in a water bath maintained at 37°C. Sections were briefly rinsed in distilled water, dehydrated through a graded series of alcohols, cleared in two changes of xylene, and coverslipped with permanent mounting medium prior to imaging. Images were captured using an Olympus SZX10 microscope at 0.63x magnification. Nissl staining revealed visible brain lesions in all but two animals in the stroke group; these animals were therefore excluded from all subsequent analyses. Examples of fixed brains and Nissl-stained sections are shown in Figure 1.

**Figure 1.**
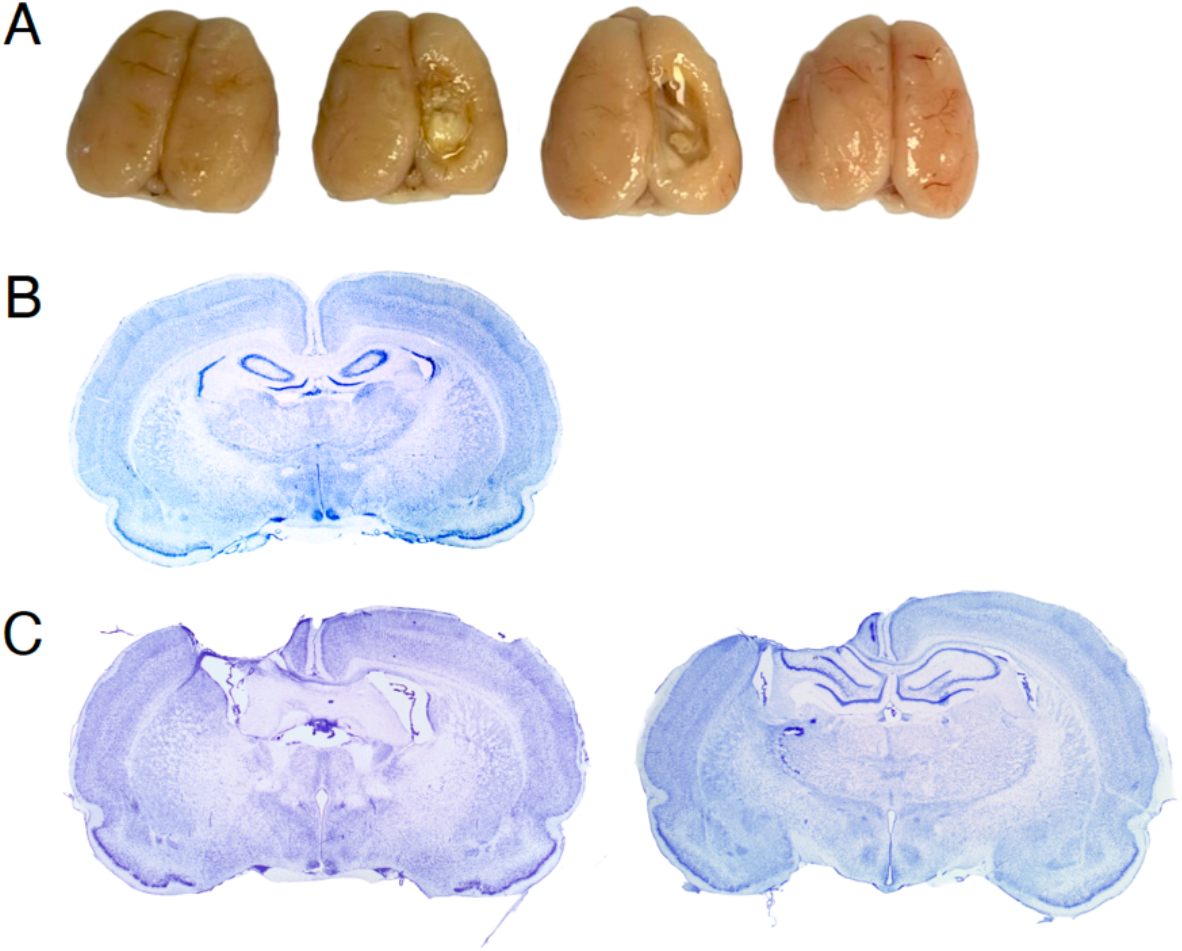
Representative brains and Nissl-stained sections following photothrombotic stroke. (A) Dorsal views of representative brains fixed in neutral buffered formalin (NBF) following removal of the cerebellum. From left to right: sham, stroke with right hemisphere lesion, stroke with right hemisphere lesion, sham. (B) Representative Nissl-stained coronal brain section from a sham animal. (C) Nissl-stained coronal sections from a stroke animal with a left hemisphere lesion, showing two different anterior-posterior levels from the same brain. The colour balance, brightness, and contrast of the images in B and C were adjusted in ImageJ.

### Quantification of sarcomere length and dispersion, fascicle length, and serial sarcomere number

Both hindlimbs from each animal were dissected by severing the pelvis bone to keep the femur intact. All muscles and other tissues were dissected from the bone, except for the extensor digitorum longus (EDL) and lateral gastrocnemius (LG) muscles. We pinned the limbs to the base of a silicone-lined container with the knee and ankle positioned at 90° angles. Once secured, the container was filled with 10% NBF to fully submerge the tissue, and the limbs were fixed for approximately 5 days. Following fixation, we rinsed the limbs in 1x phosphate-buffered saline (PBS), and the EDL and LG were carefully removed with a scalpel and placed in fresh PBS for 2 hours. The muscles were then immersed in 30% nitric acid for 9–12 hours to digest the connective tissue, rinsed in PBS, and transferred to glycerol for 2 weeks. After 2 weeks, fascicle bundles were teased from the whole muscle using a needle under a microscope. Because fascicle lengths vary across a given muscle, particularly in the architecturally-complex LG, we ensured fascicles were selected from the same location and compartment across muscles and animals. Fascicle bundles were then mounted onto slides, with five slides of fascicle bundles prepared per muscle and maintained in glycerol. We imaged the fascicle bundles using an Olympus SZX10 microscope.

Fascicle and sarcomere lengths were analyzed using custom-written code in Mathematica (Wolfram Research, Champaign, IL, USA). To measure fascicle length from the images, we first calibrated each image using the ruler included in the image. For each fascicle bundle, we manually digitized three fascicles per bundle by placing 30–50 points along the length of each fascicle. We used the digitized coordinates to generate a cubic spline representing the trajectory of each fascicle and calculated fascicle length by numerically integrating the arc length of the spline. We then converted the measured lengths from pixels to millimetres using the image-specific calibration factor and averaged the lengths of the three fascicles to obtain a single fascicle length value for each fascicle bundle.

We quantified sarcomere length using laser diffraction (ter Keurs et al., 1978). We placed the slides horizontally above a white surface and aligned the samples so that the laser beam was perpendicular to the slide. The laser was directed at individual fascicle bundles, producing a diffraction pattern on the white surface below, which we imaged with a ruler in frame for spatial calibration. For each fascicle bundle, we acquired diffraction patterns at five locations distributed along its length, including near each end and at three approximately equally spaced locations between them. To analyze the diffraction patterns, we calibrated each image using the ruler and calculated the mean pixel intensity across each column to generate an intensity profile. We then identified the central (zeroth-order) and first-order diffraction peaks and determined their positions from the cumulative intensity distribution, defining the centre of each peak as the position corresponding to 50% of its cumulative area. We calculated median sarcomere length from the spacing between the zeroth and first-order peaks using the diffraction grating equation, which relates sarcomere length to laser wavelength and diffraction angle. The diffraction angle was calculated from the peak spacing and the distance between the fascicle and projection surface. We calculated sarcomere length using both the left and right first-order peaks relative to the central peak and averaged these values to account for slight laser misalignment. We calculated sarcomere length dispersion as the distance between the positions corresponding to 25% and 75% of the cumulative area of each first-order peak and averaged the values from the left and right peaks for each sampling location. Finally, we averaged the measurements from the five sampling locations to obtain a single median sarcomere length and sarcomere length dispersion value for each fascicle bundle.

### Statistical analysis

We assessed beam traversal performance using a linear mixed-effects model implemented in R with the *lme4* package and *lmer* function. Mean percentage of missteps relative to the pre-surgery baseline was modeled as the continuous response variable, with group, side, and phase of recovery (weeks 1 and 2 versus weeks 16 and 17) included as fixed effects, along with their two- and three-way interactions. Rat was included as a random intercept to account for repeated measurements within animals. Post hoc comparisons were conducted using estimated marginal means with the *emmeans* package, with Holm correction applied for multiple comparisons. We used the same linear mixed-effects modeling framework to examine sarcomere length, fascicle length, and serial sarcomere number, with separate models fit for each outcome. For these analyses, group, muscle, and side were specified as fixed effects, including their interactions, and rat was included as a random intercept. To compared body mass at the beginning of the study prior to surgery and at harvest between stroke and sham groups, we used independent-samples Welch’s t-tests. Data were assessed for normality within each group using the Shapiro-Wilk test and for homogeneity of variance using Levene’s test.

## Results

Body mass at the beginning of the study prior to stroke or sham surgery did not differ between stroke and sham groups (281.5 ± 18.6 g vs. 270.6 ± 13.0 g, respectively; p = 0.13). Similarly, body mass at sacrifice did not differ between sham and stroke groups (361.0 ± 20.7 g vs. 351.4 ± 21.5 g, respectively; p = 0.35).

The beam traversal task revealed a significant effect of group (p < 0.001), limb side (p < 0.001), and a group x limb side interaction (p < 0.001; Figure 1). Post hoc comparisons showed that stroke animals made significantly more missteps with the paretic limb than control animals during both the early (weeks 1,2; p < 0.001) and late (weeks 16,17; p < 0.001) recovery periods, whereas non-paretic limb performance did not differ between groups at either time point (p > 0.25). Within the stroke group, rats made significantly more missteps with the paretic limb than the non-paretic limb during both early and later recovery (p < 0.001), while no side-to-side differences were observed in controls (p > 0.48). There was no significant effect of recovery phase (p = 0.56), nor were any interactions involving recovery phase significant, and post hoc comparisons confirmed that misstep rates did not differ between the first two weeks (1,2) and last two weeks (16, 17) of beam testing for either limb in either group (p > 0.16). Together, these findings indicate that stroke produced a persistent, limb-specific impairment in beam traversal performance that remained evident throughout the 4.5 months after stroke.

Sarcomere length did not differ between stroke and control animals (p = 0.29) or between paretic and non-paretic limbs (p = 0.32; Figure 2A). Sarcomere length differed significantly between muscles (p < 0.001), with longer sarcomeres in the EDL than the LG (2.27 ± 0.01 vs. 2.21 ± 0.01 μm). No significant interactions between group, muscle, or limb side were observed (all p > 0.24). Sarcomere length dispersion did not differ between stroke and control animals (p = 0.63) or between paretic and non-paretic limbs (p = 0.68; Figure 2B). Dispersion differed significantly between muscles (p < 0.001), with lower dispersion in the LG than the EDL (0.10 ± 0.003 vs. 0.12 ± 0.003 μm). No significant interactions between group, muscle, or limb side were observed (all p > 0.20).

**Figure 2.**
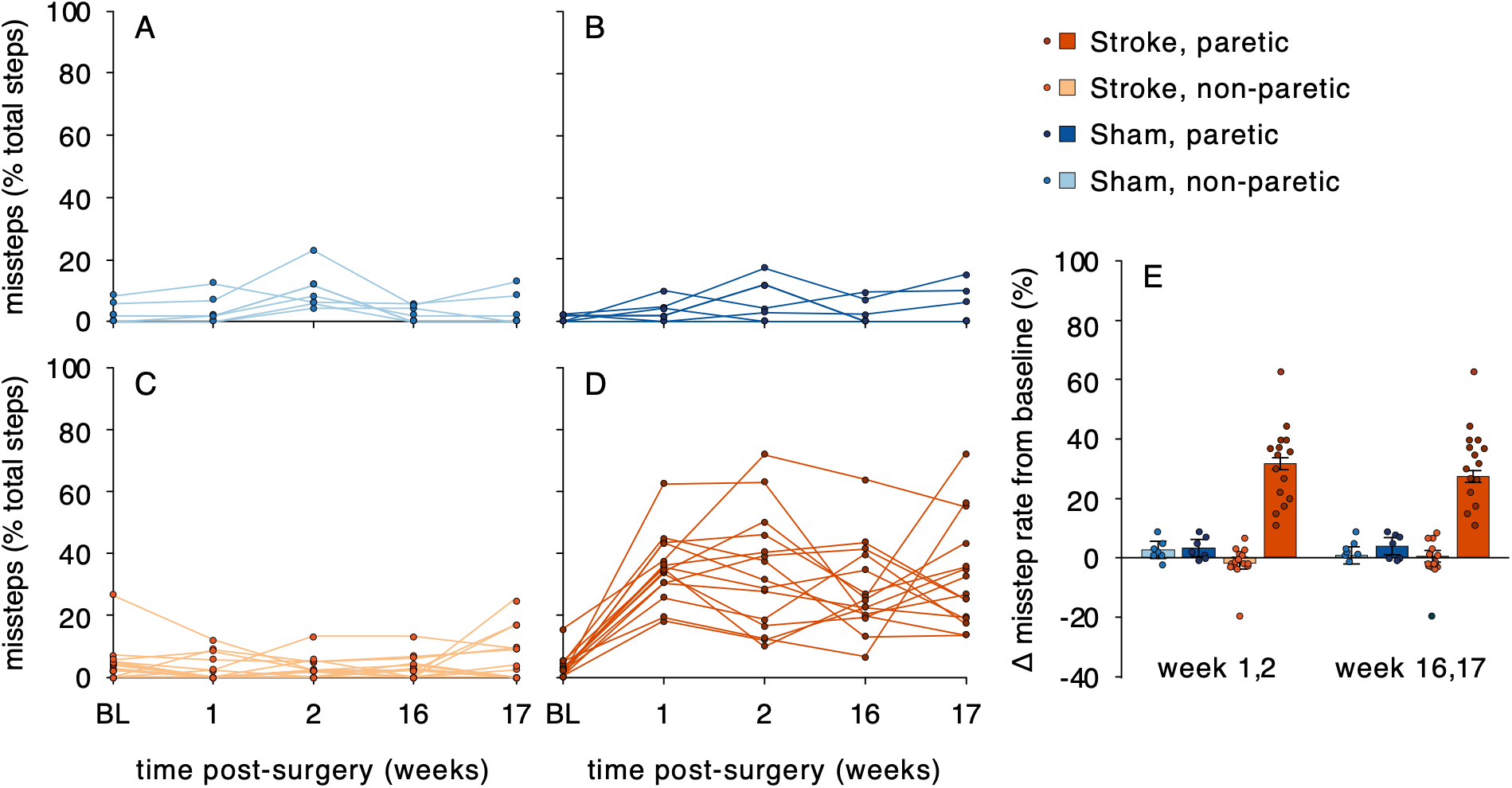
Beam traversal performance following stroke. (A–D) Missteps expressed as a percentage of total steps for the sham non-paretic (A), sham paretic (B), stroke non-paretic (C), and stroke paretic (D) limbs at baseline (BL; pre-surgery) and 1, 2, 16, and 17 weeks following surgery. (E) Change in misstep rate from baseline during the early (weeks 1,2) and late (weeks 16,17) post-surgery periods. Values were calculated by averaging the percent missteps across weeks 1 and 2 or weeks 16 and 17 and subtracting the baseline misstep rate. Bars represent group means ± SE. Colours indicate limb and surgery group: light blue, sham non-paretic; dark blue, sham paretic; light orange, stroke non-paretic; dark orange, stroke paretic.

Fascicle length differed significantly between groups and muscles (Figure 3A). Fascicles were longer in stroke animals than controls (12.6 ± 0.14 vs. 12.0 ± 0.20 mm; group main effect: p = 0.03) and longer in the LG than the EDL (13.0 ± 0.14 vs. 11.6 ± 0.14 mm; muscle main effect: p < 0.001). There was no significant effect of limb side, and no significant interactions between group, muscle, or limb side were observed (all p > 0.49).

**Figure 3.**
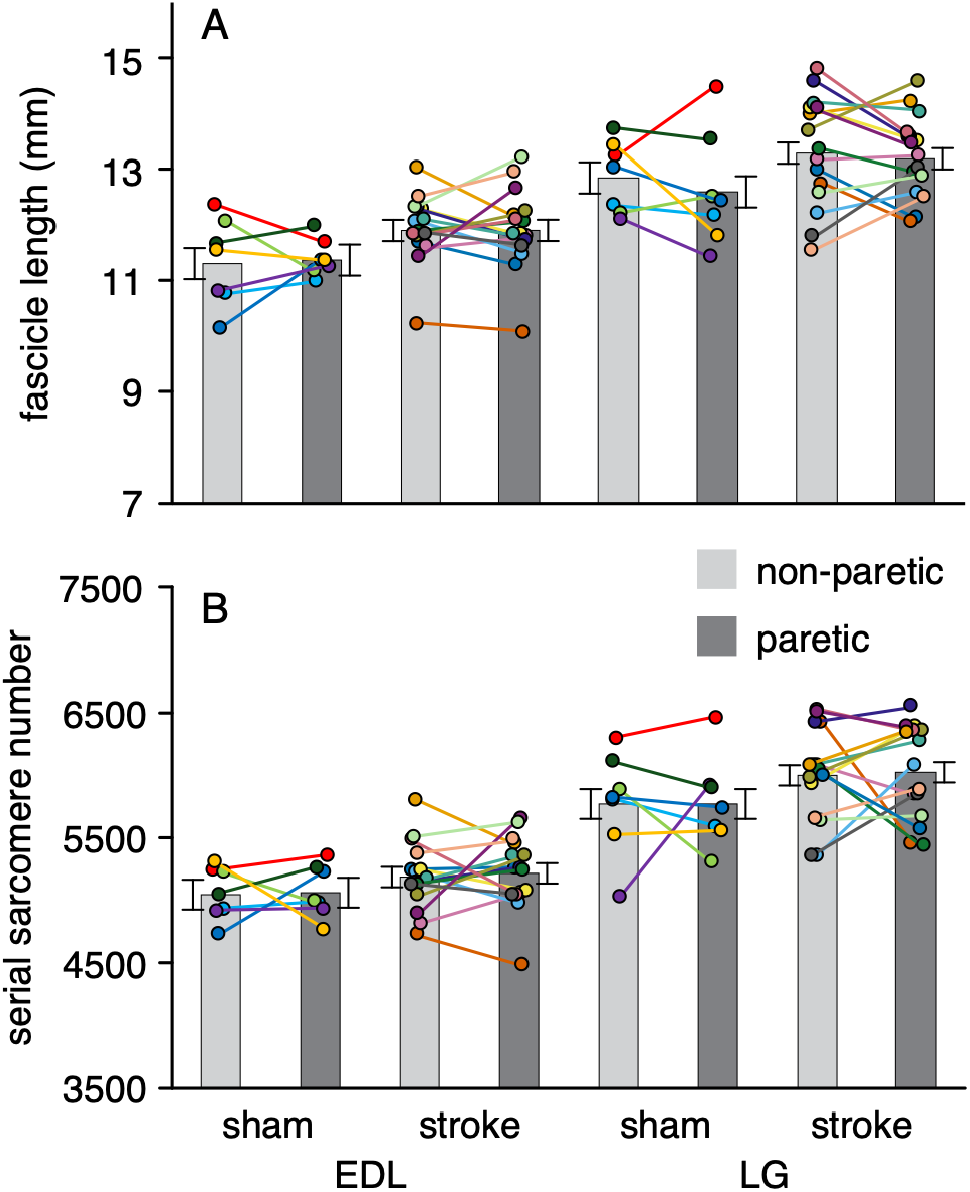
Fascicle length and serial sarcomere number in paretic and non-paretic hindlimb muscles following stroke. Fascicle length (A) and serial sarcomere number (B) are shown for the extensor digitorum longus (EDL) and lateral gastrocnemius (LG) muscles of sham and stroke animals. Within each group, light and dark gray indicate the non-paretic and paretic limbs, respectively. Lines connect measurements from the EDL and LG muscles obtained from the same animal, with each animal represented by a unique line colour. Bars represent group means ± SE.

**Figure 4.**
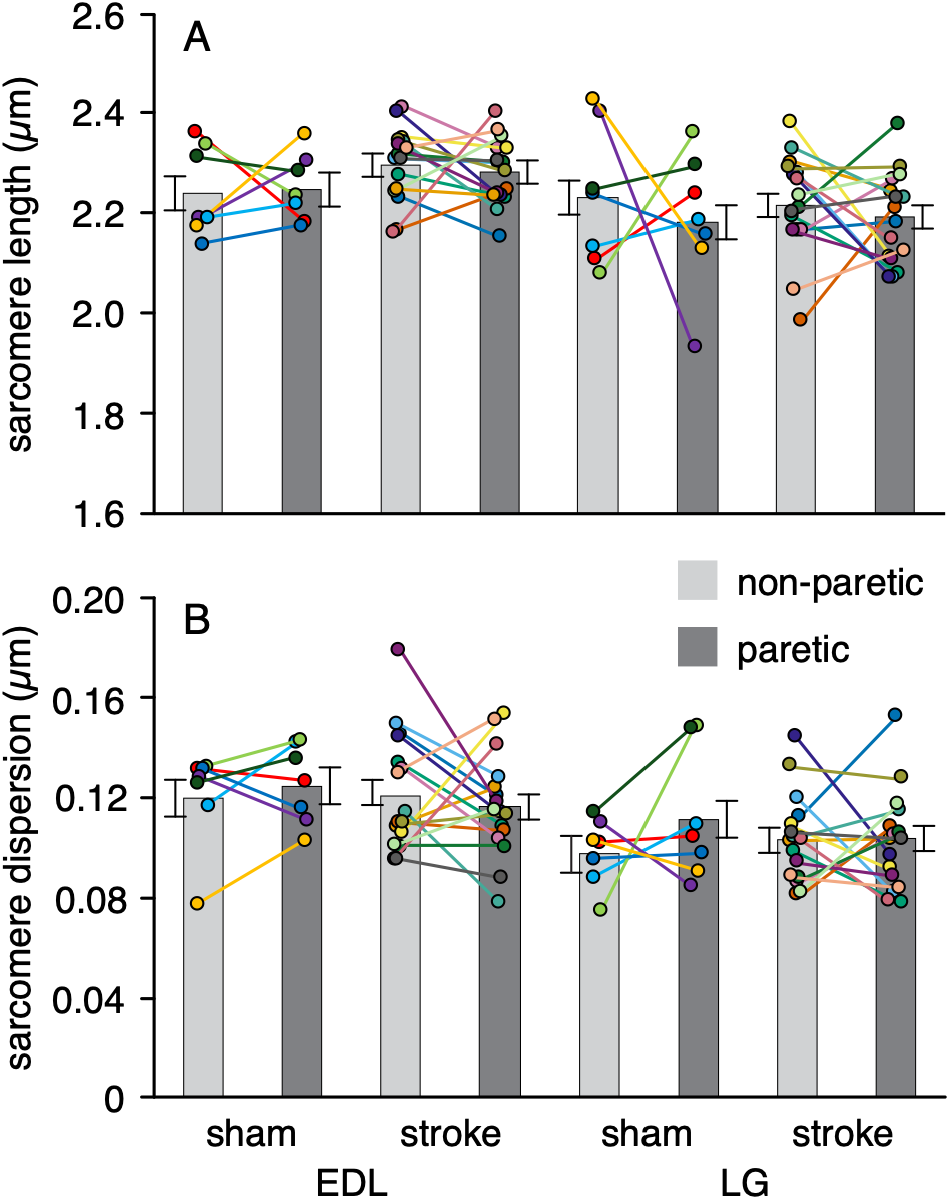
Sarcomere length and sarcomere length dispersion in paretic and non-paretic hindlimb muscles following stroke. Median sarcomere length (A) and sarcomere length dispersion (B) are shown for the extensor digitorum longus (EDL) and lateral gastrocnemius (LG) muscles of sham and stroke animals. Within each group, light and dark gray indicate the non-paretic and paretic limbs, respectively. Lines connect measurements from the EDL and LG muscles obtained from the same animal, with each animal represented by a unique line colour. Bars represent group means ± SE.

Serial sarcomere number differed significantly between muscles (p < 0.001), with more serial sarcomeres in the LG than the EDL (5889 ± 55 vs. 5128 ± 55, respectively; Figure 3B). Serial sarcomere number also differed between groups (p = 0.03), with stroke animals having more serial sarcomeres than controls (5606 ± 51 vs. 5412 ± 75, respectively). There was no significant effect of limb side (p = 0.73), and no significant interactions between group, muscle, or limb side were observed (all p > 0.47).

## Discussion

The purpose of this study was to determine whether chronic stroke alters sarcomere length, sarcomere length dispersion, fascicle length, or serial sarcomere number in paretic hindlimb muscles, and whether these changes differed between paretic and non-paretic limbs. Despite persistent motor impairments in the paretic limb during beam traversal throughout the 4.5-month recovery period, we found no effect of stroke group or limb side on sarcomere length or sarcomere length dispersion. Because motor impairment persisted throughout the post-stroke period, it is less likely that the absence of sarcomere changes reflects an initial motor deficit that was too brief to induce sarcomere adaptation or that adaptation occurred but was subsequently reversed following motor recovery. While we found no effect of limb side, we did observe a significant effect of group on fascicle length and serial sarcomere number, with stroke animals having longer fascicles and more sarcomeres in series than sham animals. Greater mechanical stimuli could theoretically promote sarcomerogenesis by chronically stretching the muscle and increasing the number of sarcomeres in series (Goldspink et al., 1974; Tabary et al., 1972). One potential mechanism for increased mechanical stimuli following stroke is increased tendon stiffness, which could result in longer muscle lengths for a given joint angle and potentially provide a stimulus for sarcomerogenesis. However, although studies examining tendon stiffness following stroke are limited, those available have found reduced stiffness in tendon on the paretic compared to non-paretic side in vivo in humans (Dias et al., 2019; Zhao et al., 2009) and in both limbs relative to age-matched controls (Dias et al., 2019). Thus, the longer fascicles and greater number of sarcomeres in series likely reflect a group difference unrelated to stroke. Because serial sarcomere number was calculated from fascicle and sarcomere length, the group difference in serial sarcomere number is consistent with the difference in fascicle length, given the absence of a significant group difference in sarcomere length. We also observed significant differences between the two muscles in fascicle length, sarcomere length, sarcomere length dispersion, and serial sarcomere number. Differences in fascicle length likely reflect known differences in the architecture of the EDL and LG muscles (Eng et al., 2008), whereas differences in sarcomere length may reflect differences in relative muscle length at the selected joint angles.

Our finding that fascicle length was unaffected by paresis in the stroke animals contrasts with the substantially shorter fascicles reported in the paretic compared with the non-paretic biceps brachii by Adkins et al. (2021) using extended field-of-view ultrasound. This discrepancy may reflect differences in the passive mechanical conditions under which fascicle length was assessed. In a study of passive human medial gastrocnemius, Gao et al. (2009) found no difference in fascicle length between paretic and non-paretic muscles when the ankle was plantarflexed with the knee at 90°, whereas paretic fascicles were significantly shorter when the ankle was dorsiflexed with the knee at the same angle. The authors proposed that the shorter fascicles at the longer muscle length reflected greater stiffness of the paretic muscle. Similarly, Kwah et al. (2012) found no difference in medial gastrocnemius fascicle length between individuals with stroke and controls at slack length, but fascicles were shorter in the stroke group when external forces were applied to lengthen the muscle. Together, these findings suggest that stroke-related differences in fascicle length may become apparent primarily when muscles experience higher passive forces, rather than reflecting differences in fascicle length at slack or low passive force. Consistent with this interpretation, diffusion tensor imaging of the medial gastrocnemius in individuals with chronic stroke found no difference in fascicle length between paretic and non-paretic limbs when muscles were assessed under conditions expected to produce relatively low passive forces (D’Souza et al., 2020a). Together, these findings suggest that stroke-related differences in fascicle length may depend on muscle length and the magnitude of passive force, which could explain differences among studies.

If greater muscle stiffness contributed to the shorter fascicle lengths in the paretic muscle in Adkins et al. (2021), shorter sarcomere lengths would also be expected at the same joint angles, yet they found no difference in sarcomere length between limbs. However, sarcomere length can be heterogeneous along a muscle (Huxley and Peachey, 1961; Lichtwark et al., 2018; Moo et al., 2016), and microendoscopy, which was used to measure sarcomere length in that study, samples only a very small region of the muscle. The technique can visualize approximately 20-50 sarcomeres concurrently (Llewellyn et al., 2008), which, relative to the approximately 40,000 sarcomeres in series reported by Adkins et al. (2021), represents only 0.05–0.13% of the total sarcomeres in series. Thus, the sampled region may not have experienced the same length change as the fascicle. It is therefore possible that the serial sarcomere number calculated from fascicle length and locally measured sarcomere length was overestimated in the paretic muscle if the sampled sarcomeres did not reflect sarcomere length throughout the fascicle. This possibility remains speculative but highlights the difficulty of relating whole fascicle length to sarcomere length when only a small number of sarcomeres from a limited region of the fascicle are sampled.

Methodological differences may have also contributed to the shorter fascicles reported by Adkins et al. (2021). Although extended field-of-view ultrasound can capture images spanning the entire muscle, it can be difficult to position the probe such that the entire length of an individual fascicle is visualized as a distinct continuous structure. More commonly, only portions of fascicles can be clearly visualized, with the remainder of the fascicle inferred based on its apparent trajectory and assumed insertion into the aponeurosis. This may be particularly challenging in paretic muscle, where potentially increased fat and fibrotic tissue could alter ultrasound image quality and contrast (Pillen et al., 2008; Pillen et al., 2009; Reimers et al., 1993). The biceps brachii may present an additional challenge because it is a fusiform muscle with fascicles that vary in length from the central region toward the outer edges, which could make it difficult to consistently select fascicles from the same anatomical region in the paretic and non-paretic limbs. Furthermore, Adkins et al. reported reduced cross-sectional area of the paretic biceps brachii, consistent with a loss of muscle fibers in parallel, which would alter the muscle architecture and make it more difficult to select fascicles from comparable regions between limbs. Together, these factors could have increased uncertainty in the fascicle length measurements and contributed to the observed difference between paretic and non-paretic limbs.

The absence of sarcomere length adaptations despite persistent motor impairment in our stroke animals suggests that altered sarcomere length may not result from abnormal neural drive alone but may require neural impairment to occur prior to or during a period of longitudinal muscle growth. During development, muscles normally increase in length through sarcomerogenesis or the addition of sarcomeres in series. In CP, this process may be impaired, resulting in fewer serial sarcomeres such that remaining sarcomeres must stretch to longer lengths to accommodate increases in muscle length as the skeleton grows (Leonard et al., 2019; Lieber and Fridén, 2002; Mathewson et al., 2015; Smith et al., 2011). In contrast, stroke occurs after the musculoskeletal system is largely developed, so the neural injury occurs after the period when muscles must undergo sarcomerogenesis to accommodate longitudinal growth. Thus, the absence of altered sarcomere length or serial sarcomere number in our chronic stroke model may reflect the adult timing of the neural injury; if stroke occurred earlier in life and was followed by substantial skeletal growth, impaired sarcomerogenesis could potentially result in fewer sarcomeres in series and overstretching of the remaining sarcomeres. Spasticity and contracture have also been proposed as potential mechanisms contributing to muscle remodelling and altered growth in CP (Handsfield et al., 2022). However, rodent stroke models are not thought to consistently reproduce the spasticity observed following human stroke (Bandela et al., 2026) and therefore may not entirely capture the neural and mechanical environment that could contribute to impaired sarcomerogenesis. Thus, the absence of differences in sarcomere length in our stroke model may reflect the adult timing of the neural injury, although differences in spasticity and other aspects of the post-stroke mechanical environment between rodent models and humans may also contribute.

An important consideration is whether the preservation of sarcomere and fascicle length observed in the present study extends to other muscles and regions of the body. We examined the EDL and LG, which have opposing actions at the ankle as dorsiflexors and plantarflexors, respectively, and observed similar effects of stroke in both muscles. However, post-stroke motor impairments can produce muscle-specific patterns of activation, including greater activation of flexor muscles in the upper limb and extensor muscles in the lower limb, as well as abnormal coupling between muscles that would normally be activated independently (Li et al., 2021). Thus, the neural and mechanical environments experienced by the EDL and LG may differ from those experienced by muscles that may be more involved in post-stroke flexor or extensor synergies. Although our finding that both a dorsiflexor and plantarflexor exhibited preserved sarcomere and fascicle length suggests that this preservation may not be specific to a single muscle action, studies of additional muscles across limbs are needed to determine whether sarcomere alterations after stroke are muscle-specific or depend on the pattern of altered neural activation.

While we found no difference in sarcomere and fascicle lengths between paretic and non-paretic sides, there may be changes in muscle structure and function at larger scales that alter force output and contribute to motor impairments. For example, studies of individuals with stroke have reported alterations in muscle size, although findings vary across studies and muscles, with reductions in muscle volume (D’Souza et al., 2020a) and cross-sectional area (Ramsay et al., 2011; Ryan et al., 2011), while others have reported no differences in cross-sectional area (D’Souza et al., 2020a). Increased intramuscular fat (D’Souza et al., 2020b) and altered fibre-type properties (Ansved et al., 1996; McDonald et al., 2021; Noguchi et al., 2023; Snow et al., 2019) have also been reported. Thus, the persistent motor impairment observed after stroke may involve muscle adaptations that were not captured by the measures examined in the present study. Examining muscle structure and function across multiple scales will be important for determining the extent to which skeletal muscle contributes to motor impairment following stroke.

Several characteristics of the experimental model may have influenced the muscle response to stroke, including the relatively young and healthy nature of the animals, their female sex, and their rapid return to spontaneous activity following stroke. Rats were 14 weeks old at the time of stroke induction and did not have the age-related comorbidities that are common among people who experience stroke (Appelros et al., 2021; Elamy et al., 2020). These factors may influence muscle health and the response to neural injury, although their relevance to sarcomere and fascicle length adaptations is less clear. Age-related changes in sarcomere length and serial sarcomere number are likely relatively modest (Hinks and Power, 2025; Power et al., 2021) compared with the substantial changes produced by chronic alterations in muscle length, such as immobilization (Goldspink et al., 1974; Tabary et al., 1972). All animals in the present study were female because male rats grow substantially larger over the study and would become too large to reliably perform the beam traversal task by the end of the post-stroke period. As with age and comorbidities, sex may influence other aspects of muscle biology (Emmert et al., 2024;

Rosa-Caldwell and Greene, 2019); however, there is limited evidence that it substantially influences sarcomerogenesis, suggesting that our findings likely apply to males as well. In addition, the animals resumed walking and general movement within approximately 1 hour of surgery, indicating that they experienced relatively minimal sustained reduction in mobility during the acute post-stroke period. This rapid return to mobility may differ from humans because rats are quadrupedal and can better compensate for paresis using their other three limbs, and humans recovering from stroke may spend substantial time inactive following stroke while in the clinical setting (Bernhardt et al., 2004; Chen et al., 2020). However, despite their early return to spontaneous activity, the animals exhibited persistent motor impairments throughout the post-stroke period, suggesting that alterations in neural input and muscle loading likely remained despite early mobilization. Thus, while early mobilization, young age, absence of comorbidities, and female sex may influence other aspects of muscle adaptation following stroke, these factors likely had minimal effect on the sarcomere and fascicle results in the present study.

Overall, the findings of this study indicate that chronic stroke does not necessarily result in alterations in sarcomere length, sarcomere length dispersion, fascicle length, or serial sarcomere number in adult skeletal muscle, despite persistent motor impairment. The preservation of sarcomere structure in both paretic and non-paretic limbs suggests that persistent changes in neural drive following stroke may not, by themselves, be sufficient to produce the sarcomere changes observed in some other neurological conditions. Instead, altered sarcomerogenesis may depend on additional factors, including the timing of neural injury relative to longitudinal muscle growth and the duration and mechanical consequences of altered muscle use. The persistent motor deficits observed in the absence of changes in sarcomere or fascicle length also suggest that stroke-related muscle adaptations may occur at other structural or functional scales that were not captured in the present study. Together, these findings highlight the importance of considering the timing, severity, and context of neural injury, as well as muscle-specific and species-specific factors, when interpreting skeletal muscle adaptations following stroke.

## Acknowledgements

We thank Hannah Smith for assistance with beam traversal training and Nicole Conquergood for assistance with tissue harvest. We also thank Dr. Richard Dyck for providing guidance with surgical procedures and the veterinary and husbandry staff at the University of Calgary’s Life and Environmental Sciences Animal Research Centre for their support with animal care.

## Author Contributions

Conceptualization: S.A.R.; Methodology: S.A.R., S.N.D., D.C.; Investigation: S.A.R., S.N.D., R.S.; Formal analysis: S.A.R.; Visualization: S.A.R.; Supervision: S.A.R., T.R.L., W.H.; Project administration: S.A.R., T.R.L.; Funding acquisition: S.A.R., W.H.; Writing – original draft: S.A.R.; Writing – review & editing: S.A.R., S.N.D., T.R.L., R.S., D.C., W.H.

